## Supplementary figures and tables for "Orthogonal inducible control of Cas13 circuits enables programmable RNA regulation in mammalian cells"

Supplementary Figure 1

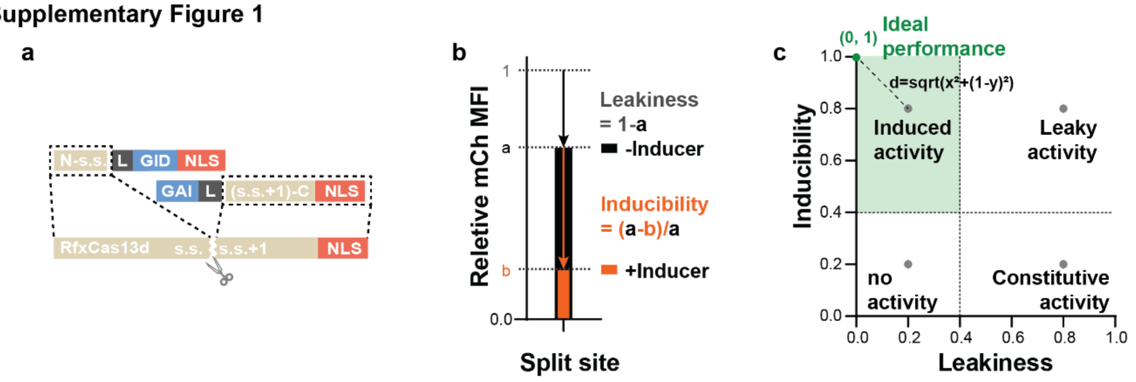


**Supplementary Figure 1**

A. Construct design of GA-induced split RfxCas13d for initial screening. The design of the N-terminal moiety, from N- to C-terminus, is the Cas13d fragment from the N-terminus to the split site (s.s.) (tan) followed by a GS-linker (black), a GID domain (blue), and an NLS tag (orange). The design of the C-terminal moiety, from N- to C-terminus, is the GAI domain (blue), a GS-linker (black), and then the Cas13 fragment from the ss to the C-terminus (tan), and an NLS tag (orange).

B. Relative mCherry MFI is the processed measurement of Cas13 activity in screening experiments. The MFI of mCherry of each sample is normalized to that of iRFP (within the same sample) to account for knockdown specificity and transfection variations across samples. Then these normalized values are normalized to a negative control transfected with only mCherry and iRFP without the Cas13 system. When Cas13 is not active, the relative mCherry MFI should be 1, and it should be less than 1 when it is active and achieves mCherry-specific knockdown. Leakiness is defined as the difference in the relative mCherry MFI between the uninduced condition and 1. The inducibility of a screened system is defined as the fraction difference in relative mCherry MFI between GA versus vehicle control-induced groups.

C. This leakiness versus inducibility plot has four quadrants of activity. Inducible split designs that show high inducibility and low leakiness are located in the upper left quadrant. Designs with high leakiness and high inducibility are located in the upper right quadrant. The bottom left quadrant represents inactive designs. The bottom right quadrant represents designs with constitutive activity (WTCas13 will be located here). The ideal performance of an inducible split Cas13 design is at (0,1), and the distance to the ideal performance is used to linearly evaluate system performance in CID orientation screening experiments.

Supplementary Figure 2

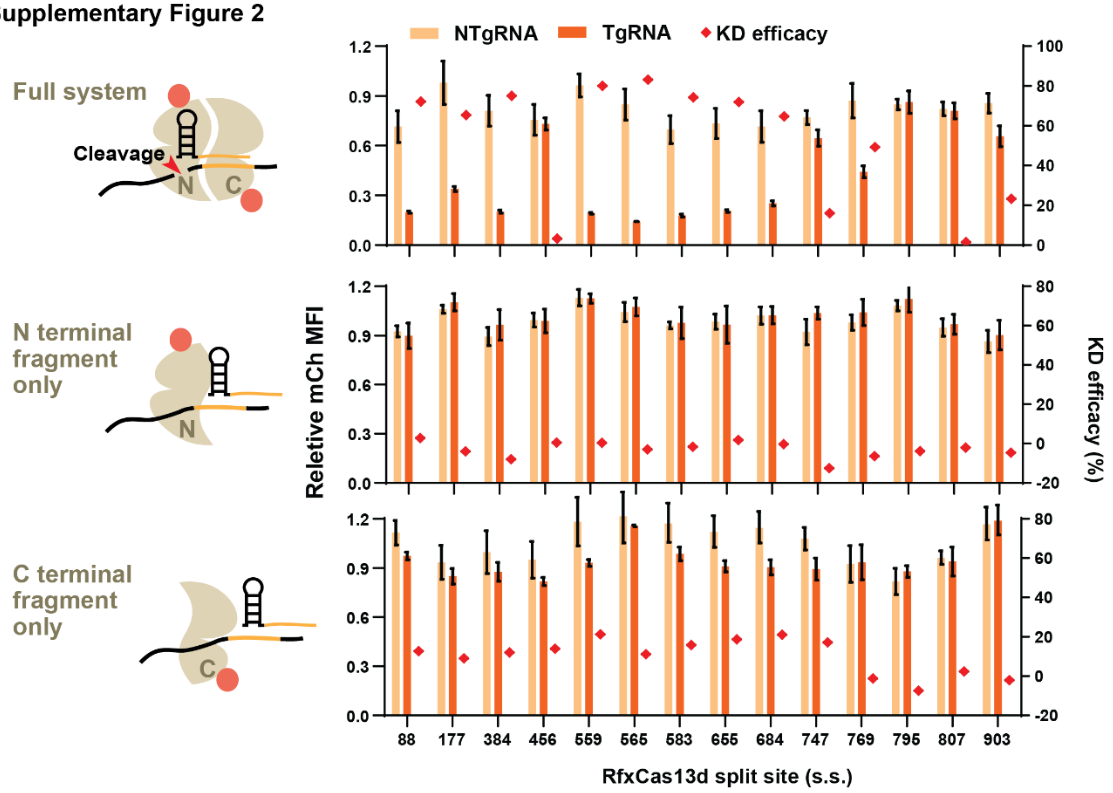


**Supplementary Figure 2**

Split Cas13s without recruitment domain are transfected in pairs, only an N- or a C-terminal fragments with either an NT gRNA or mCherry targeting gRNA. Split sites that showed great leakiness (88/89, 177/178, 384/385, 559/560, 583/584, 655/656, and 684/685) in initial screening could reconstitute and become active for mCherry-specific knockdown in the presence of targeting gRNA without recruitment domains, while split sites offering no activity in initial screening remained inactive without recruitment domains. No unpaired N- or C-terminal fragment could knock down mCherry with targeting gRNA.

**Supplementary Figure 3**

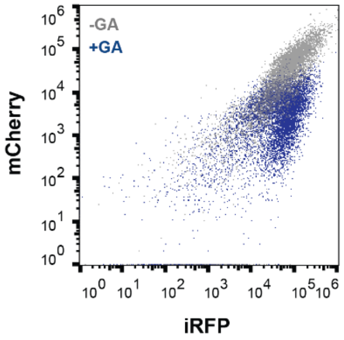


**Supplementary Figure 3**

Flow cytometry data for mCherry and iRFP expression in the presence of the design N507-GAI-NLS/GID-508C-NLS without (grey) and with GA (navy) induction. mCherry was specifically knockdown down, demonstrated by the downward shift of the blue population with respect to the grey. Although cells with lower transfection efficiency showed insufficient mCherry knockdown, the relative MFI of the entire live population was knocked down 80% with GA induction.

**Supplementary Figure 4**

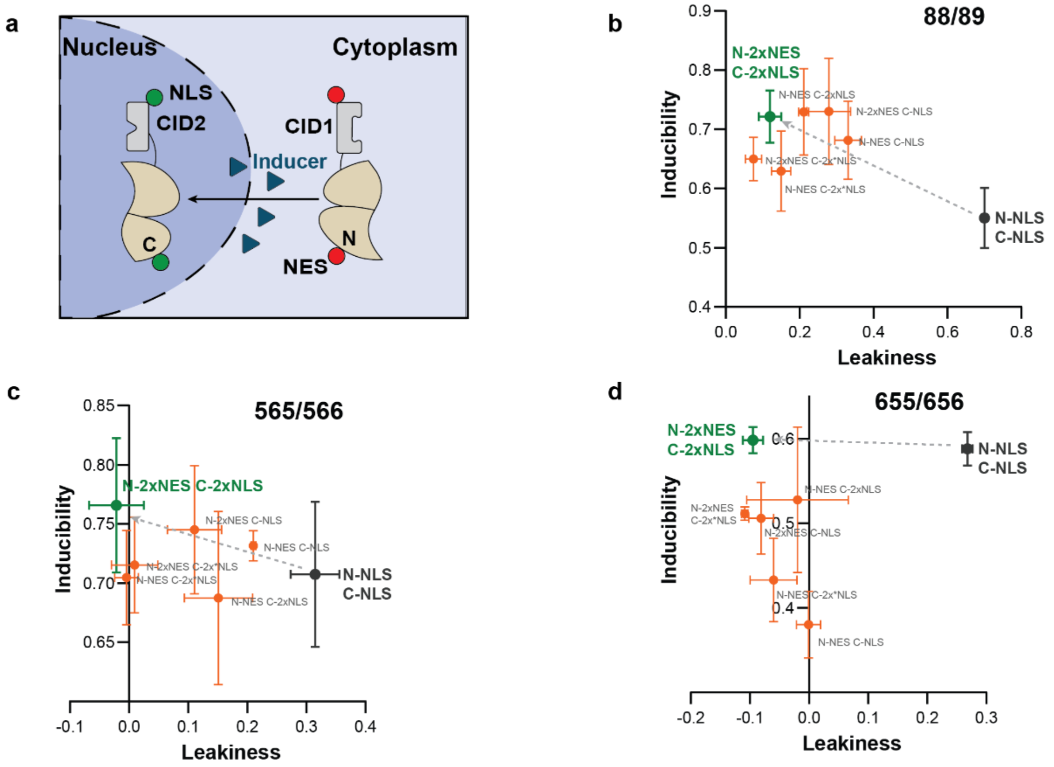


**Supplementary Figure 4**

A. A schematic showing the balanced shuttling of Cas13 splits between the cytosol and nucleus. In the absence of an inducer, the N- and C- terminal moieties are separately sequestered in the cytosol or the nucleus preventing them from spontaneous dimerization. With the inducer added, N and C split moieties would bind together, thus offering stronger ligand-responsive ON activity of Cas13 and better dynamic range between ON and OFF states.

B. A leakiness of 0.7 by the GA-inducible RfxCas13d split sites 88/89 was rescued with enhanced inducibility. The best performance was offered by the design NES-N88-GID-NES/NLS-GAI-89C-NLS.

C. A leakiness of 0.31 by the GA-inducible RfxCas13d split site 565/566 was rescued without a negative impact on inducibility. The best performance was offered by the design NES-N565-GID-NES/NLS-GAI-566C-NLS.

D. A leakiness of 0.28 by the GA-inducible RfxCas13d split sites 655/656 was rescued without a negative impact on inducibility. The best performance was offered by the design NES-N655-GID-NES/NLS-GAI-656C-NLS.

Supplementary Figure 5

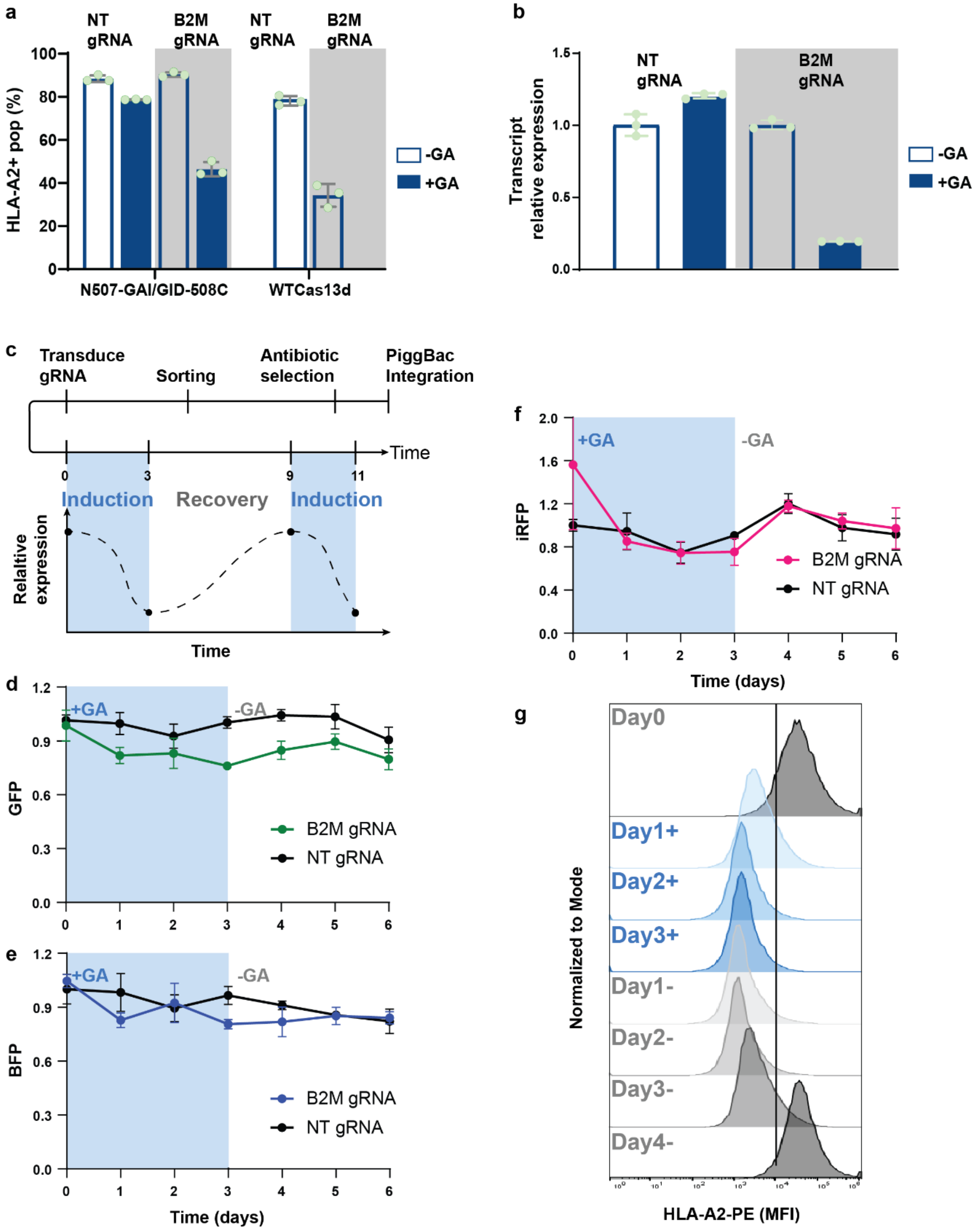


**Supplementary Figure 5**

A. B2M targeting gRNA was first tested with transient transfection. GA induced downregulated HLA surface expression in 50% transfected cells, while GA with non-targeting gRNA or the targeting gRNA without GA conditions did not reduce HLA expression. GA induced knockdown efficiency was comparable to that by WT RfxCas13.

B. In a stable cell line highly expressing the GA induced split Cas13d effector, transiently transfected gRNA led to >80% knockdown of the B2M transcript upon GA induction, while GA or the targeting gRNA alone led to no targeted knockdown of B2M transcripts.

C. Experiment timeline for the reversibility of GA induced split Cas13d activity. HEK293FT cells were transfected with the PiggyBac transposon system containing an expression cassette for GA induced split Cas13. After transfection, the cell line was selected with antibiotics for at least 3 passages. Next, cell sorting was performed to select for clones with strong Cas13 expression. The sorted cell line was subsequently transduced with lentivirus carrying the gRNA to generate a stable cell line equipped with the full B2M targeting GA induced split Cas13d system. B2M expression of stable cells experiencing GA induction and recovery was measured as surface HLA expression using flow cytometry.

D. Bystander fluorescent protein expression (GFP) was not disturbed during GA induction and withdrawal. GFP expression cassette was delivered with gRNA using lentivirus.

E. BFP, which was expressed with N-terminal Cas13d, also remains constant throughout the induction.

F. iRFP was expressed with C-terminal split moiety. The expression of all system components remains constant throughout the reversibility experiment.

G. Histograms showing HLA surface staining distribution of cell population stably expressing B2M targeting GA induced split Cas13d system. After 2 days of GA induction, the histogram shows a full population shift to a low HLA-expressing state. 2 days after recovery from GA induction, the population shifts to the right expressing more HLA with full recovery from GA induced HLA knockdown on day 4 after GA withdrawal.

Supplementary Figure 6

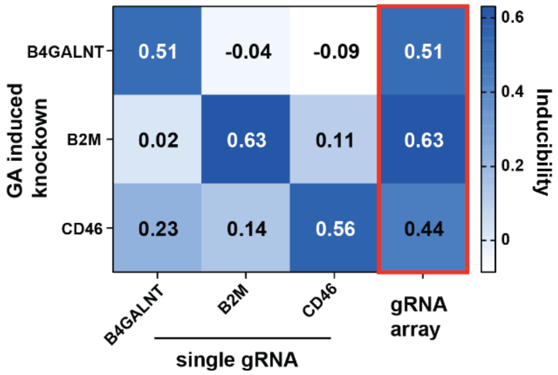


**Supplementary Figure 6**

Heat map showing induced knockdown of 3 endogenous targets-B4GALNT, B2M, and CD46, with either single target mature gRNA or a gRNA array. Every single gRNA is orthogonal to other targets as each transcript is only inducibly knocked down with its corresponding single gRNA. The gRNA array successfully knocked down all 3 targets simultaneously with inducibility comparable to their single mature gRNAs.

**Supplementary Figure 7**

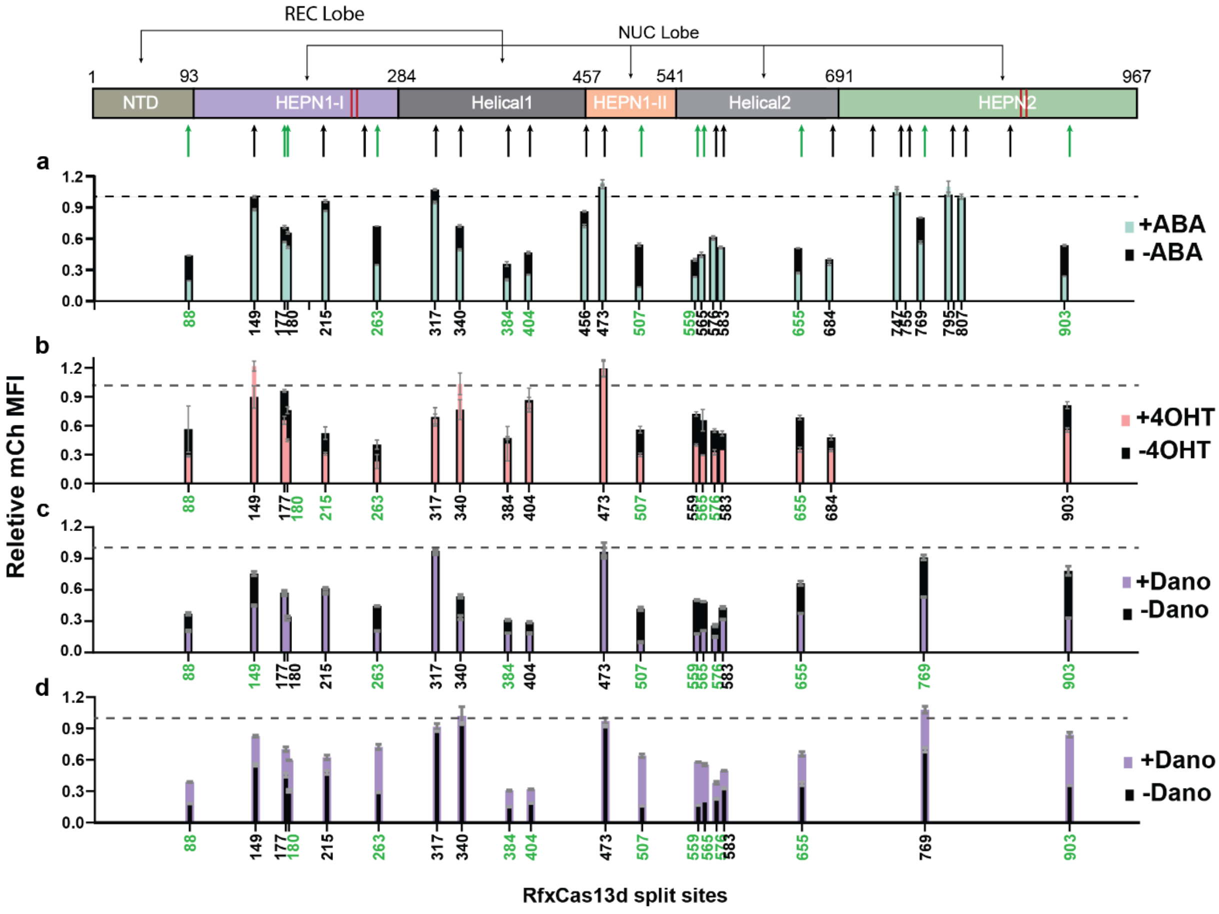


**Supplementary Figure 7**

Screening results of split RfxCas13d with the ABA, 4-OHT, Dano induced dimerization domains, and Dano inhibited dimerization domains. Green arrows are split Cas13d pairs that generated >0.4 inducibility with the GA induced dimerization system in the initial screening. Split 88/89, 263/264, 507/508, and 655/656 consistently generated >0.4 inducibility with all inducible systems.

A. The system design (which denotes which CID fragment is connected to split Cas13 and the location of the NLS) for ABA induced split Cas13d design is N-ABI-NLS/PYL-C-NLS. ABA induced dimerization system generally offered lower inducibility (black portion of each bar) compared to GA induced dimerization system, and fewer splits generated >0.4 inducibility. Split 177/178, 180/181, and 565/566, when induced with ABA, were not as efficient at mCherry knockdown, leading to their reduced inducibility. Split 903/904 offered a stronger induced knockdown and >0.4 inducibility.

B. The design for a 4-OHT induced split Cas13d used in screening is N/ERT2-C-NLS. 6/8 split sites with >0.4 GA responsive inducibility generated >0.4 inducibility with the 4-OHT-inducible dimerization system. Additionally, split 215/216 and 576/577 generated high inducibility with 4-OHT induced dimerization system.

C. With the Dano induced dimerization system, the design used in the screening is N-dNS3-NLS/DNCR-C-NLS. Split 177/178 and 180/181 did not generate >0.4 inducibility due to substantial leaky activity in the uninduced condition. 576/577, 769/770, and 903/904 generated higher inducibility than when they were coupled with the GA system.

D. With the Dano inhibited dimerization system, the design used in the screening is as N-dNS3-NLS/ANR-C-NLS. 11 out of 19 screened split sites generated >0.4 inducibility with the Dano inhibited dimerization system. However, compared to other split sites that generated >0.4 with at least 1 other previous system, the unique split sites 384/385 and 404/405 were highly leaky and highly active in uninduced conditions.

Supplementary Figure 8

a

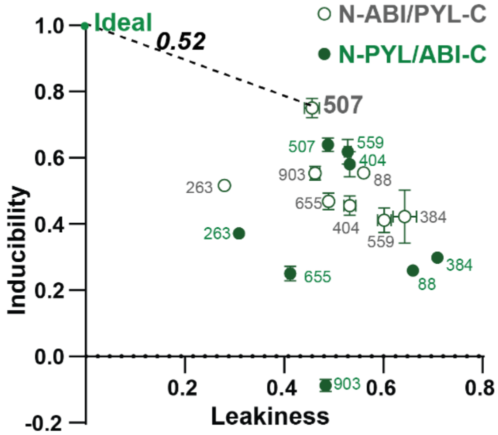


| Distance to Ideal Performance | | |
|---|---|---|
| N-ABI/PYL-C | s.s. | N-ABI/PYL-C |
| 0.72 | 88 | 0.99 |
| 0.56 | 263 | 0.70 |
| 0.86 | 384 | 1.00 |
| 0.76 | 404 | 0.68 |
| 0.52 | 507 | 0.61 |
| 0.84 | 559 | 0.65 |
| 0.72 | 655 | 0.86 |
| 0.64 | 903 | 1.19 |
| *Leakiness/Inducibility* | | |
| 0.46/0.75 | | 0.49/0.64 |

b

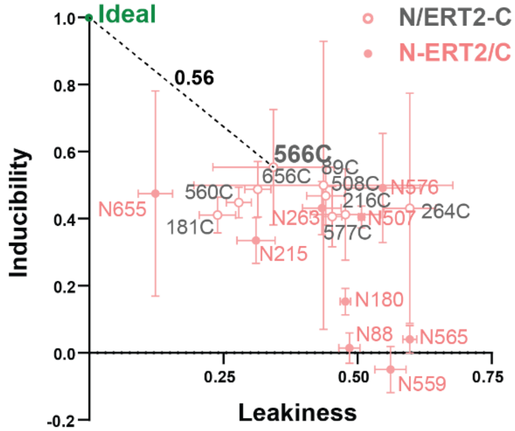


| Distance to Ideal Performance | | |
|---|---|---|
| N/C-ERT2 | s.s. | N-ERT2/C |
| 0.66 | 88 | 1.10 |
| 0.62 | 559 | 1.19 |
| 0.56 | 565 | 1.13 |
| 0.60 | 655 | 0.54 |
| 0.64 | 180 | 0.97 |
| 0.76 | 215 | 0.73 |
| 0.82 | 263 | 0.71 |
| 0.69 | 507 | 0.78 |
| 0.75 | 576 | 0.75 |
| *Leakiness/Inducibility* | | |
| 0.34/0.55 | | 0.12/0.48 |

c

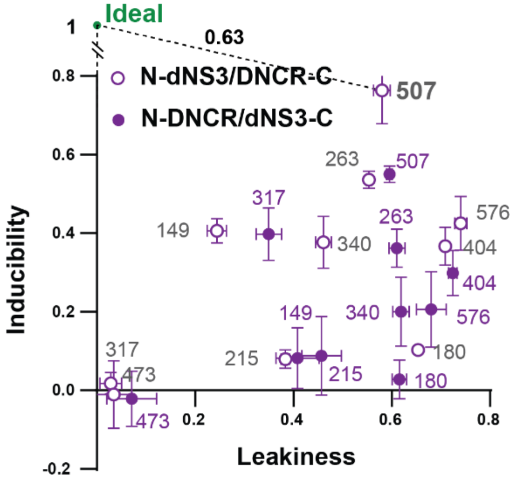


| Distance to Ideal Performance | | |
|---|---|---|
| N-dNS3 | s.s. | N-DNCR |
| 0.64 | 149 | 1.00 |
| 1.11 | 180 | 1.15 |
| 1.00 | 215 | 1.02 |
| 0.72 | 263 | 0.88 |
| 0.98 | 317 | 0.70 |
| 0.77 | 340 | 1.01 |
| 0.95 | 404 | 1.01 |
| 1.01 | 473 | 1.02 |
| 0.63 | 507 | 0.75 |
| 0.94 | 576 | 1.05 |
| *Leakiness/Inducibility* | | |
| 0.58/0.76 | | |

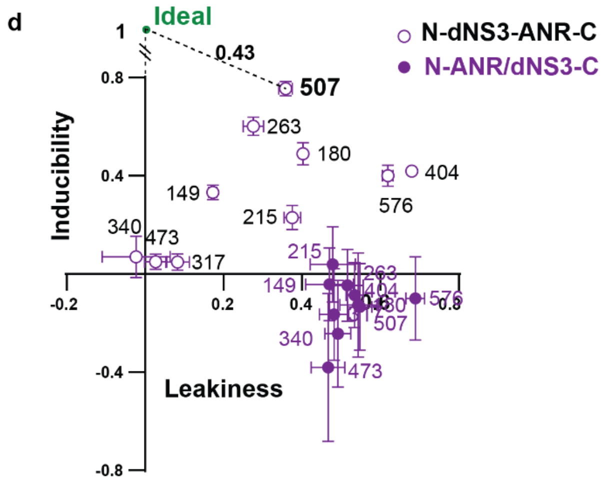


| Distance to Ideal Performance | | |
|---|---|---|
| N-dNS3 | s.s. | N-ANR |
| 0.69 | **149** | 1.14 |
| 0.65 | **180** | 1.25 |
| 0.86 | **215** | 1.07 |
| 0.48 | **263** | 1.17 |
| 0.96 | **317** | 1.26 |
| 0.93 | **340** | 1.34 |
| 0.89 | **404** | 1.21 |
| 0.95 | **473** | 1.46 |
| 0.43 | **507** | 1.26 |
| 0.86 | **576** | 1.30 |
| ***Leakiness/Inducibility*** | | |
| 0.36/0.76 | | |

**Supplementary Figure 8**

A. Comparison between two different orientations for ABA induced split Cas13d design (>0.4 inducibility) (N-ABI/PYL-C and N-PYL/ABI-C). The N-PYL/ABI-C orientation generally led to reduced inducibility across splits compared N-ABI/PYL-C. The orientation N-ABI/PYL-C generated better performance with more screened split sites. The best design was N507-ABI/PYL-508C, with a distance of 0.52 to the ideal, 0.46 leakiness, and 0.75 inducibility.

B. Comparison between two different orientations for 4-OHT induced split Cas13d design (>0.4 inducibility) (N/C-ERT2 and N-ERT2/C). The N-ERT2/C orientation generally led to higher leakiness and reduced inducibility across splits. The orientation N/C-ERT2 generated better performance with more screened split sites. The best design was N565/566C-ERT2, with a distance of 0.56 to the ideal, 0.34 leakiness, and 0.55 inducibility.

C. Comparison between two different orientations for Dano induced split Cas13d design (>0.4 inducibility) (N-dNS3/DNCR-C and N-DNCR/dNS3-C). The N-DNCR/dNS3-C orientation generally led to reduced inducibility across splits. The orientation N-dNS3/DNCR-C generated better performance with more screened split sites. The best design was N507-dNS3/DNCR-508C, with a distance of 0.63 to the ideal, 0.58 leakiness, and 0.76 inducibility.

D. Comparison between two different orientations for Dano induced split Cas13d design (>0.4 inducibility) (N-dNS3/ANR-C to N-ANR/dNS3-C). The N-ANR/dNS3-C orientation generally led to reduced inducibility across splits. The orientation N-dNS3/ANR-C generated better performance with more screened split sites. The best design was N507-dNS3/ANR-508C, with a distance of 0.43 to the ideal, 0.36 leakiness, and 0.76 inducibility.

### Supplementary Figure 9

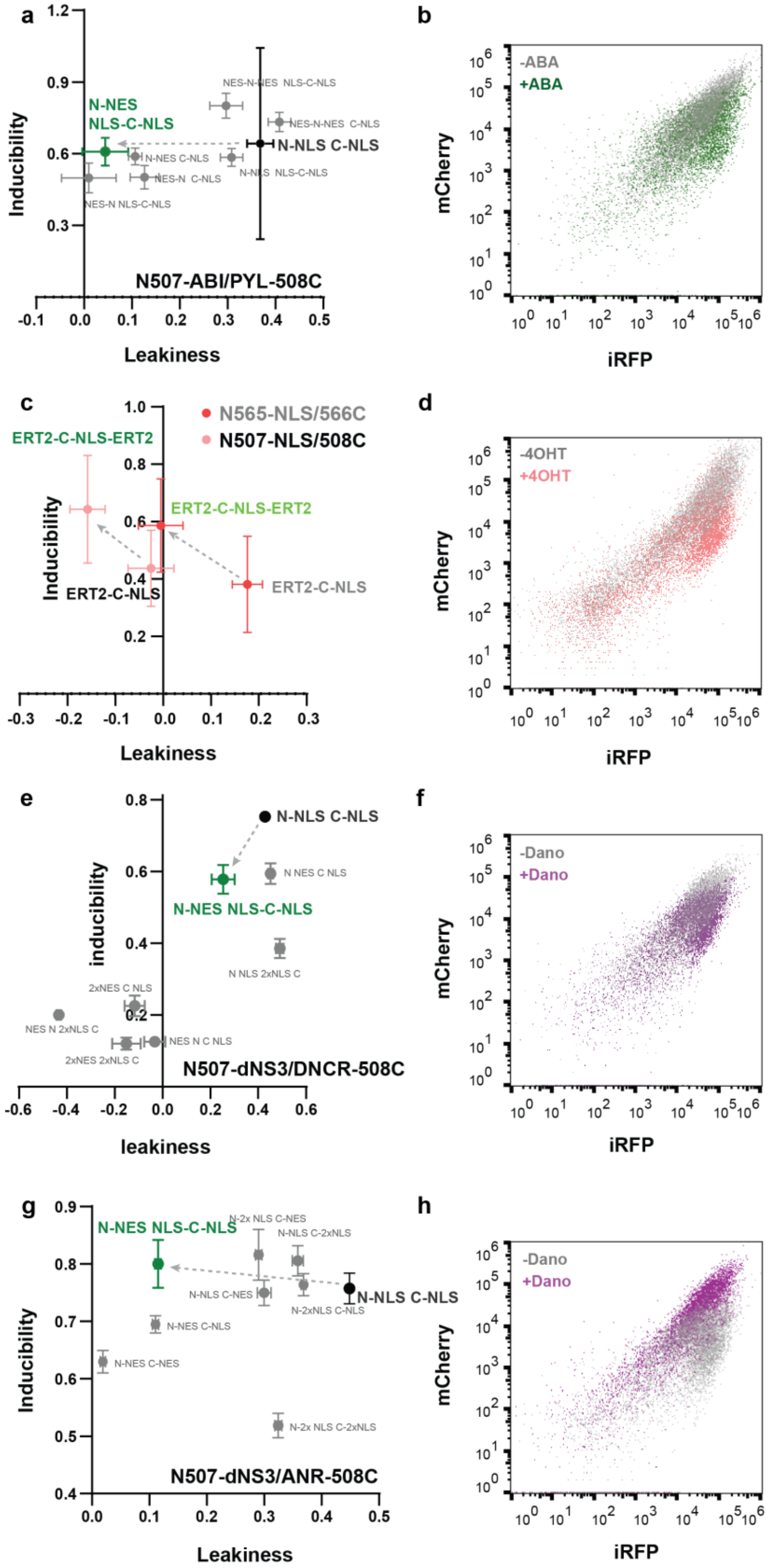

**Supplementary Figure 9**

A. NES/NLS optimization for N507-ABI/PYL-508C. N507-ABI-NES/NLS-PYL-508C-NLS fully rescued leakiness by N507-ABI-NLS/PYL-508C-NLS from 0.36 to 0.034, without reducing inducibility.

B. Flow cytometry data for mCherry and iRFP expression in the presence of the design N507-ABI-NES/NLS-PYL-508C-NLS without (grey) and with ABA (green) induction. mCherry was specifically knockdown down, demonstrated by the shift down of the green population with respect to the grey.

C. NES/NLS optimization for N565/ERT2-566C and N507/ERT2-508C. N565-NLS/ERT2-566C-NLS-ERT2 fully rescued leakiness by N565-NLS/ERT2-566C-ERT2-NLS from 0.18 to 0, enhancing inducibility from 0.39 to 0.60. N507-NLS/ERT2-508C-NLS-ERT2 showed an enhanced inducibility of 0.70 compared to other designs.

D. Flow cytometry data for mCherry and iRFP expression in the presence of the design N507-NLS /ERT2-508C-NLS-ERT2 without (grey) and with 4OHT (salmon) induction. mCherry was specifically knockdown down, demonstrated by the shift down of the salmon population with respect to the grey.

E. NES/NLS optimization for N507-dNS3/DNCR-508C. N507-dNS3-NES/NLS-DNCR-508C-NLS rescued leakiness by N507-dNS3-NLS/DNCR-508C-NLS from 0.39 to 0.21, while also reducing inducibility from 0.76 to 0.59.

F. Flow cytometry data for mCherry and iRFP expression in the presence of the design N507-dNS3-NES/NLS-DNCR-508C-NLS without (grey) and with Dano (purple) induction. MCherry was specifically knockdown down by Dano induction, demonstrated by the shift down of the purple population with respect to the grey.

G. NES/NLS optimization for N507-dNS3/ANR-508C. N507-dNS3-NES/NLS-ANR-508C-NLS rescued leakiness by N507-dNS3-NLS/ANR-508C-NLS from 0.45 to 0.12, without reducing inducibility.

H. Flow cytometry data for mCherry and iRFP expression in the presence of the design N507- N507-dNS3-NES/NLS-ANR-508C-NLS without (grey) and with Dano (purple) induction. mCherry was specifically knockdown down in the absence of Dano, demonstrated by the shift down of the grey population with respect to the purple.

Supplementary Figure 10

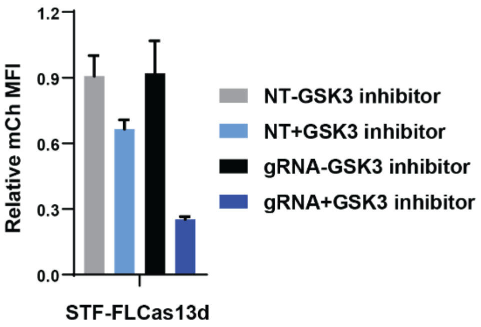


**Supplementary Figure 10**
Using the WNT-responsive SuperTOP Flash promoter to drive WTCas13d expression, 72% knockdown was achieved in response to GSK3 inhibitor induction with mCherry targeting gRNA. GSK induced mCherry knockdown in the absence of mCherry targeting gRNA.

Supplementary Figure 11

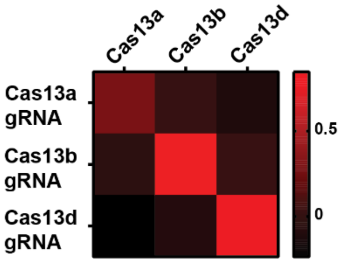


**Supplementary Figure 11**
Heat map showing orthogonality of each Cas13 ortholog- LwCas13a, PspCas13b, and RfxCas13d- with their cognate gRNAs. In addition, WT PspCas13b and WT RfxCas13d generated a more robust mCherry knockdown (~80% knockdown) than WT LwCas13a (40% knockdown).

**Supplementary Figure 12**

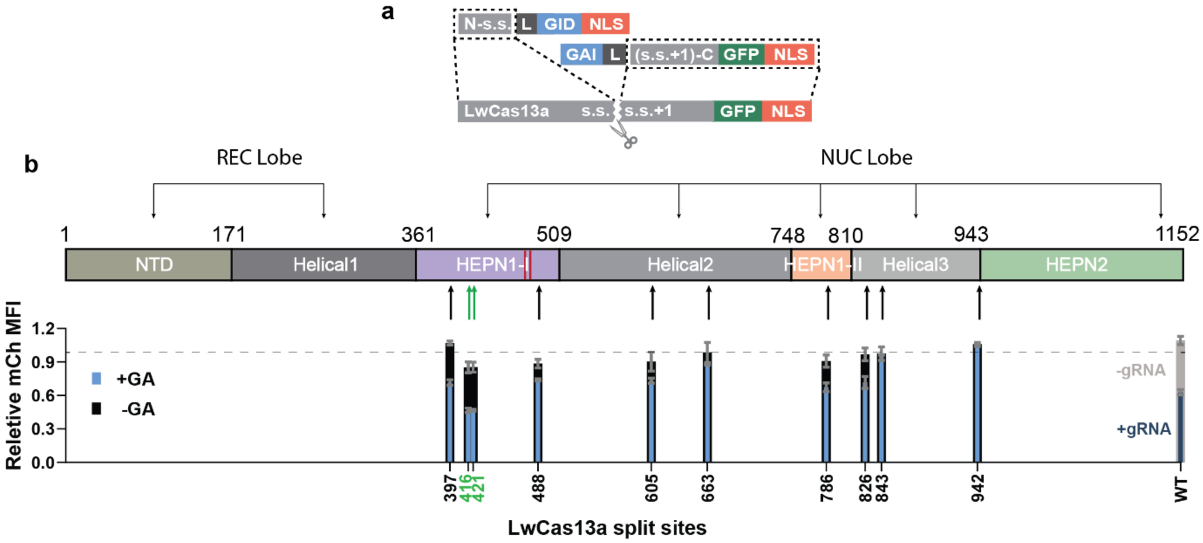


### Supplementary Figure 12

A. Design of GA induced split LwCas13a. As LwCas13 was found to be stably expression in mammalian cells when it is fused with a GFP on the C-terminus, split LwCas13 C-terminal fragments retained the fused GFP[1].

B. 10 split sites (s.s.) were screened in the GA-inducible design, and 2 generated >0.4 inducibility which is comparable to the gRNA induced mCherry knockdown by WT LwCas13a that was only 0.4. The black bar is mCherry expression without inducer, and the blue bar is lowered mCherry expression with gibberellic acid introduced.

**Supplementary Figure 13**

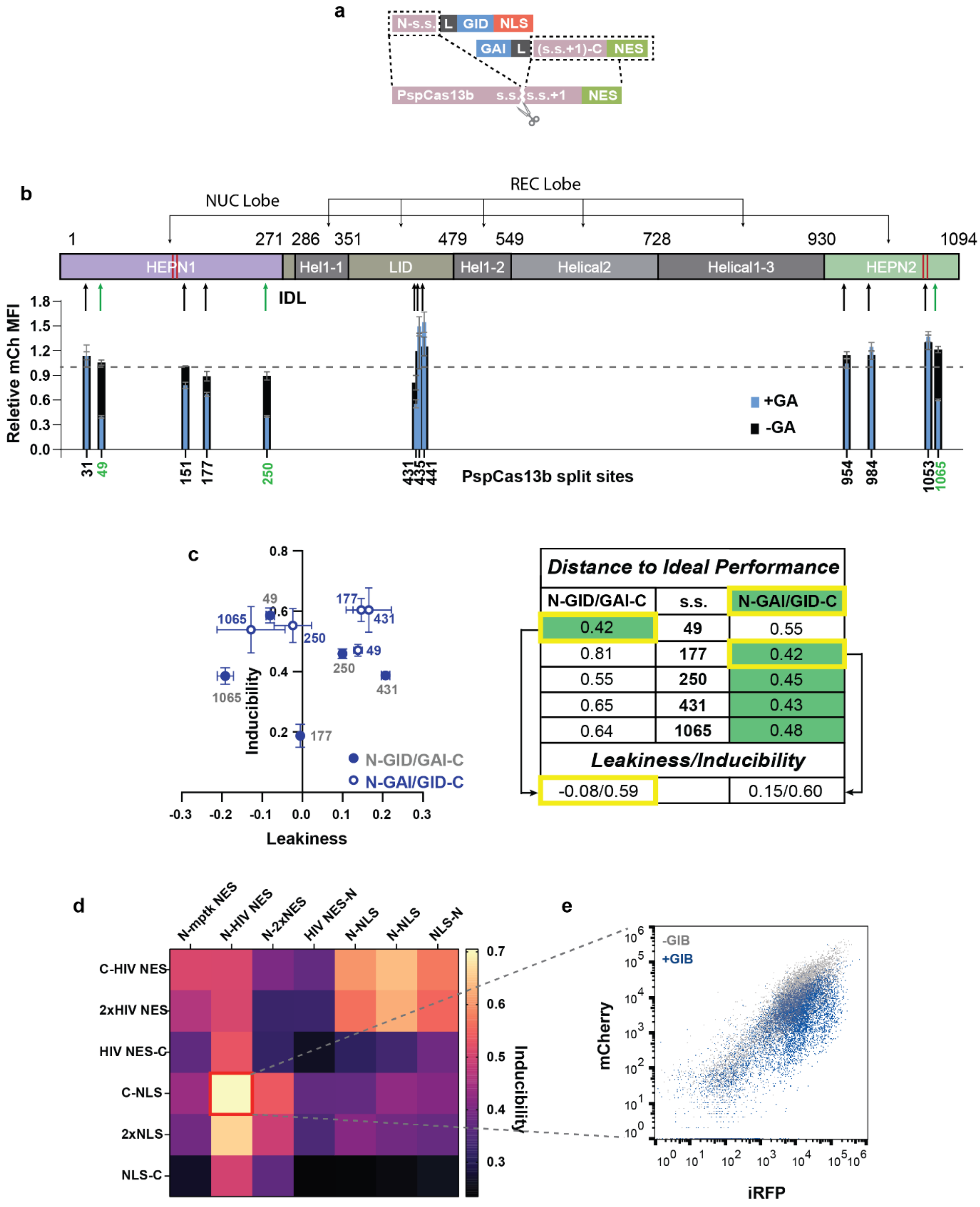


**Supplementary Figure 13**

A. Design of GA induced split PspCas13b. As PspCas13b was found to be the most efficient at targeting RNA cleavage when it is targeted to the cytosol with HIV-NES, we retained the HIV-NES tag on its C-terminus[2].

B. 3 out of 12 split sites were screened throughout the sequence of PspCas13b generated >0.4 inducibility with relatively minimal leakiness compared to what was observed in the initial RfxCas13d split screening.

C. Comparison between two different orientations for GA induced split Cas13b design from (N-GID/GAI-C and N-GAI/GID-C). The N-GAI/GID-C orientation led to increased inducibility across 4 out of 5 splits. The orientation N-GAI/GID-C generated better performance with more screened split sites. The best performance was offered by the design N49-GAI/GID-50C, with a distance of 0.42 to the ideal, minimal leakiness, and 0.59 inducibility.

D. Heatmap of inducibility generated by different designs of NES and NLS tagging for N49-GAI/GID-50C. The highest inducibility was ~0.7 offered by the design N49-GAI-NES/GID-50C-NLS.

E. Flow cytometry data for mCherry and iRFP expression in the presence of the design N49-GID-NES/GAI-50C-NLS without (grey) and with GA (navy) induction. MCherry was specifically knockdown down, demonstrated by the shift down of the navy population with respect to the grey. The relative MFI of the entire transfected population was knocked down 70% with GA induction.

**Supplementary Figure 14**

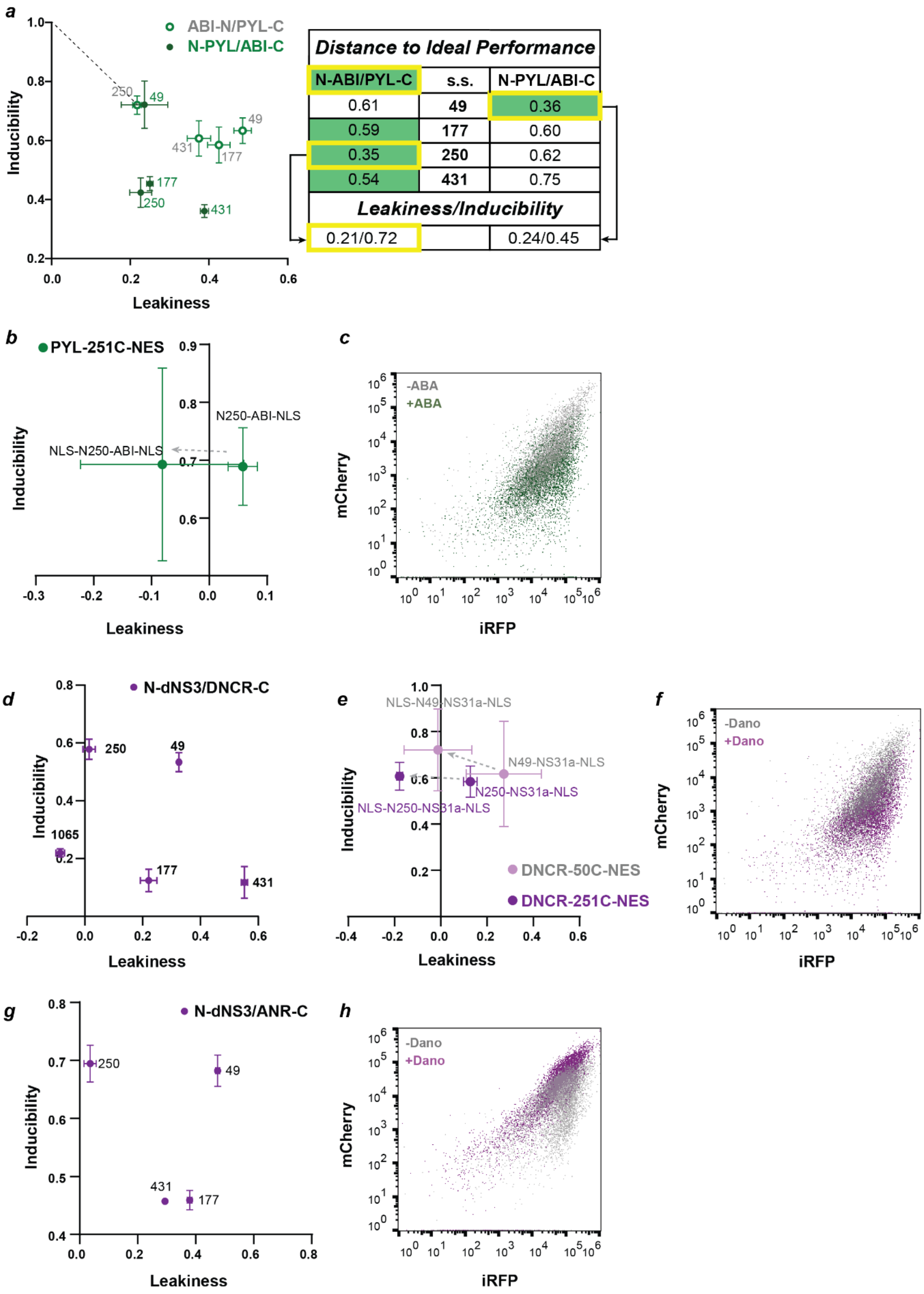

**Supplementary Figure 14**

A. Comparison between two different orientations for ABA induced split Cas13b design (N-ABI/PYL-C to N-PYL/ABI-C). The N-PYL/ABI-C reversed orientation led to increased leakiness across 3 out of 4 splits. The orientation N-ABI/PYL-C generated better performance with more screened split sites. The best performance was offered by the design N250-ABI/PYL-251C, with a distance of 0.35 to the ideal, 0.21 leakiness, and 0.72 inducibility.

B. NLS-N250-ABI-NLS/PYL-251C-NES minimized leakiness generated by design N250-ABI-NLS/PYL-251C-NES without reducing inducibility.

C. Flow cytometry data for mCherry and iRFP expression in the presence of the design NLS-N250-ABI-NLS/PYL-251C-NES without (grey) and with ABA (green) induction. mCherry was specifically knockdown down by ABA induction, demonstrated by the shift down of the green population with respect to the grey.

D. From our experience designing Dano induced split Cas13d systems, the orientation N-dNS3/DNCR-C generated higher inducibility consistently across split sites. We screened PspCas13b splits with this orientation. And split 250/251 generated ~0.6 Dano responsive inducibility with minimal leakiness.

E. Although NLS-N250-dNS3-NLS/DNCR-251C-NES did minimize the leakiness generated by N250-dNS3-NLS/DNCR-251C-NES, inducibility was also retained to be 0.6. For a system with higher inducibility, we optimized the design N49-dNS3/DNCR-50C which yielded inducibility similar to split 250/251 and additionally 0.32 leakiness. And NLS-N49-dNS3-NLS/DNCR-50C-NES did minimize the leakiness generated by N250-dNS3-NLS/DNCR-251C-NES and brought inducibility up to 0.72.

F. Flow cytometry data for mCherry and iRFP expression in the presence of the design NLS-N49-dNS3-NLS/DNCR-50C-NES without (grey) and with Dano (purple) induction. mCherry was specifically knockdown down by Dano, demonstrated by the shift down of the purple population with respect to the grey.

G. From our experience designing Dano inhibited split Cas13d systems, the orientation N-dNS3/ANR-C generated higher inducibility consistently across split sites. We screened PspCas13b splits with this orientation. And split 250/251 generated 0.70 Dano responsive inducibility with minimal leakiness.

H. Flow cytometry data for mCherry and iRFP expression in the presence of the design N250-dNS3-NLS/ANR-251C-NES without (grey) and with (purple) Dano induction. mCherry was specifically knockdown down in the absence of Dano, demonstrated by the shift down of the prey population with respect to the purple.

**Supplementary Figure 15**

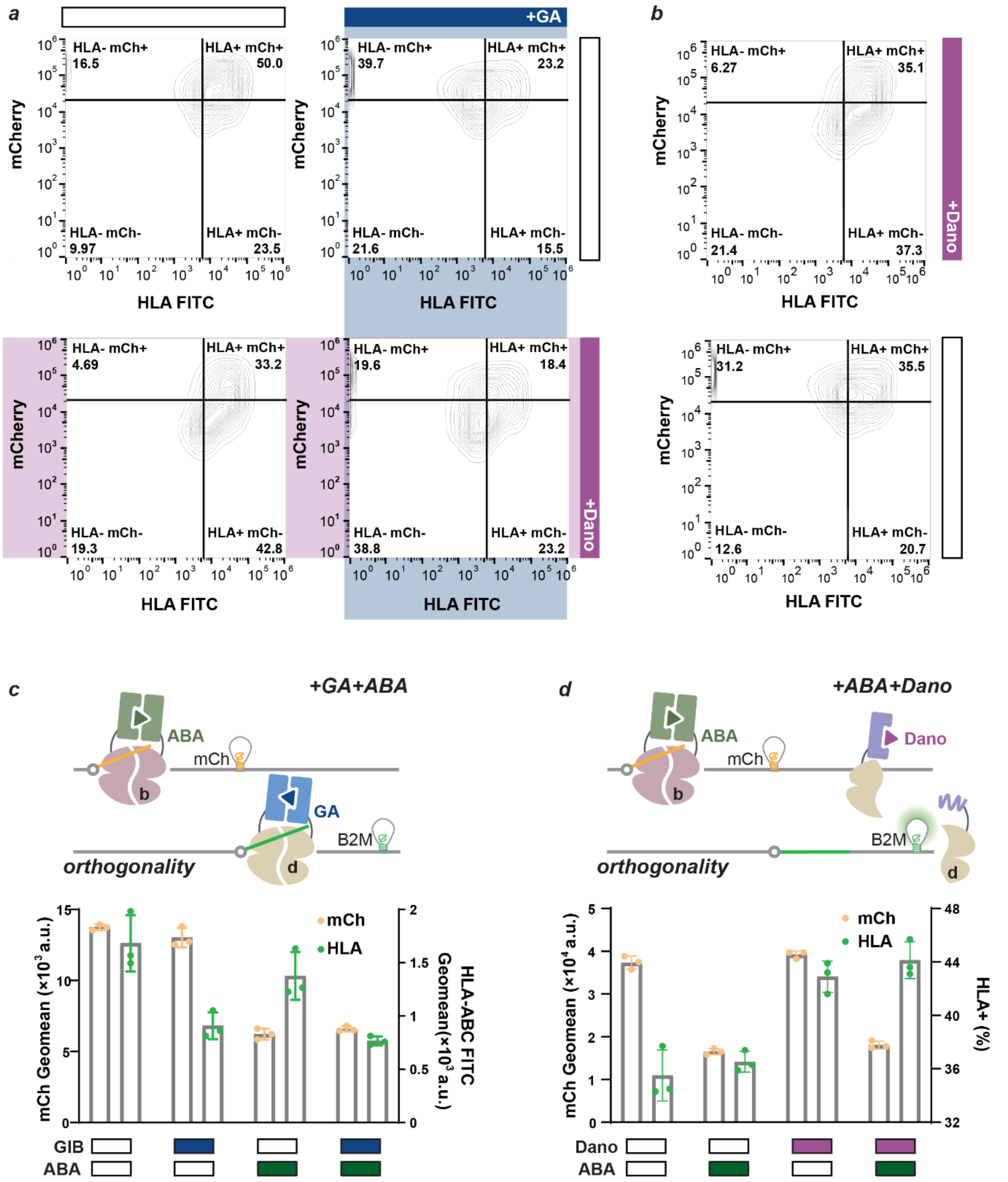

**Supplementary Figure 15**

A. Flow cytometry contour plot for the mCherry and HLA expression in cells simultaneously transfected with the mCherry-targeting Dano induced Cas13b system and the B2M-targeting GA induced Cas13d system. In the absence of Dano and GA (upper left panel), both mCherry and HLA were highly expressed; with GA induction (upper right panel), the population shifted to the left due to the reduced HLA expression; with Dano induction (bottom left panel), the population shifted downward due to the mCh specific knockdown; with dual induction of Dano and GA (bottom right panel), the population shifted to the left bottom quadrant due to the reduced expression of both mCherry and HLA.

B. Flow cytometry contour plot for the mCherry and HLA expression in cells simultaneously transfected with the mCherry-targeting Dano induced Cas13b system and the B2M-targeting Dano inhibited Cas13d system. In the absence of Dano (bottom panel), mCherry was highly expressed while HLA expression was down. With Dano induction (upper panel), mCherry was knocked down, while HLA expression was strong.

C. Orthogonal regulation of RfxCas13d and PspCas13b, respectively by GA and ABA for targeted knockdown of B2M and mCherry. B2M expression was measured as the HLA surface expression. HLA expression is decreased in the presence of GA, while mCherry expression is decreased only in the presence of ABA.

D. Orthogonal regulation of RfxCas13d and PspCas13b, respectively by Dano and ABA for targeted knockdown of B2M and mCherry. As Dano inhibited split RfxCas13d is applied, HLA expression was knocked down constitutively. Repression of HLA expression is lifted with Dano induction. mCherry expression is decreased only in the presence of ABA.

**Supplementary Figure 16**

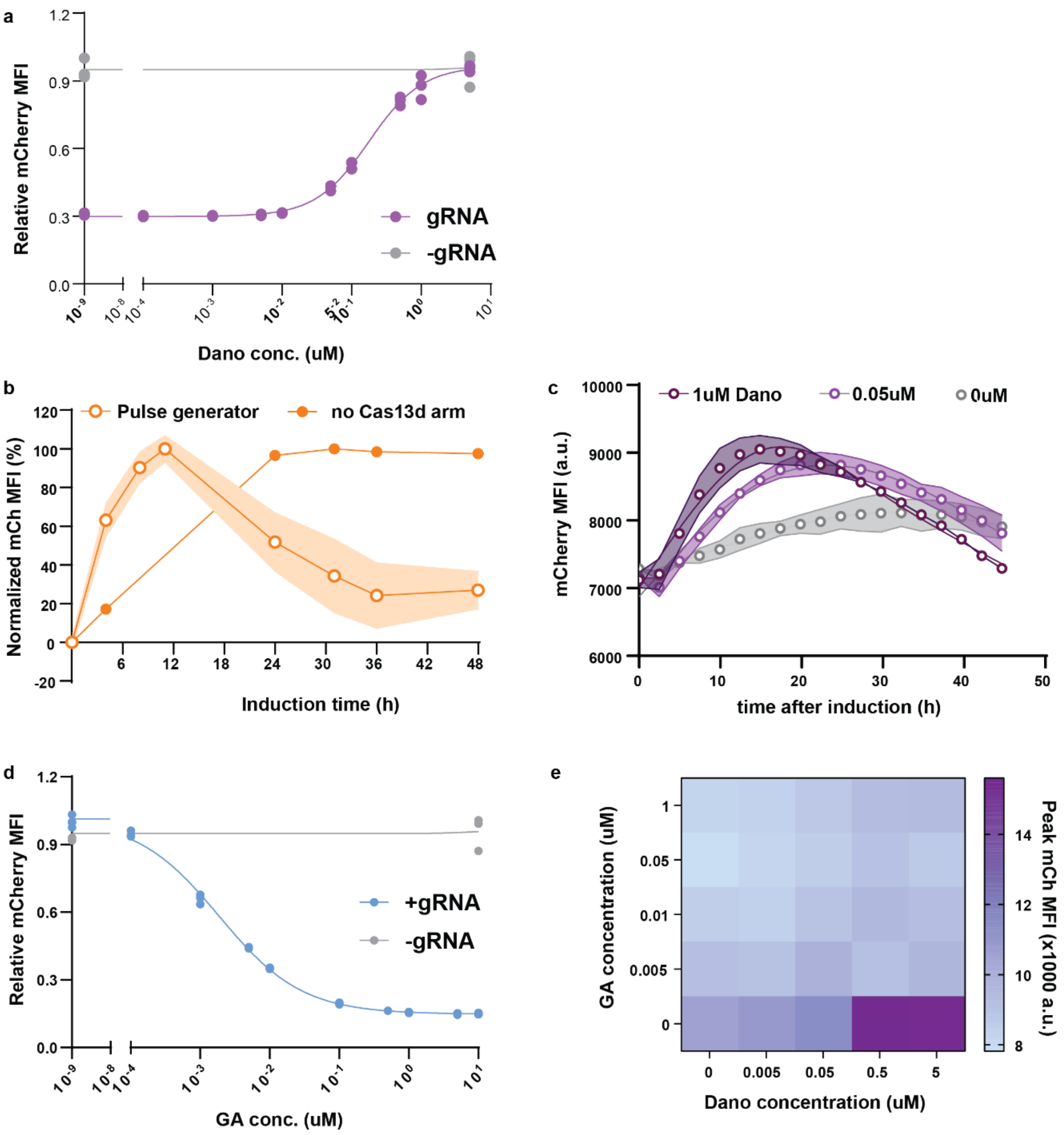


**Supplementary Figure 16**

A. Dose responsive mCherry knockdown by the Dano inhibited the Cas13b system with and without mCherry targeting gRNA. IC50=0.18uM, R squared=0.9955

B. Output expression generated by the full IFFL or IFFL without the repression arm (Cas13d) with different duration of Dano induction. Cells were transfected with either the complete IFFL circuit components or without Cas13d as the repression arm, and induced at different time points, collected, and ran flow cytometry at 48h after the first induction. Output adaptation is shown only in samples transfected with complete circuits with induction time, starting between 12 to 24h of induction and reaching a plateau at 36h.

C. mCherry MFI from real-time microscopic imaging of live cells transfected with the complete IFFL circuits and induced with 5uM, 0.05uM, and 0uM Dano. High Dano concentration results in larger peak amplitude, while no Dano induction led to only basal level mCherry expression.

D. Dose responsive mCherry expression in cells transfected with the GA induced split RfxCas13d with mCherry targeting gRNA (blue) and non-targeting gRNA (grey). IC50=0.00188uM R squared=0.9978.

E. Heatmap of the output peak amplitude generated by the dual-regulated IFFL circuit. Output expression is the strongest with high Dano concentration and no GA, which is basically the IFFL without repression arm. With GA induction, thus an active repression arm, peak amplitude is higher with higher Dano concentration and lower GA concentration.

**Supplementary Table 1. GA induced RfxCas13d system performance of designs: N-GID/GAI-C and N-GAI/GID-C.** N-GAI/GID-C is the prefered orientation for split RfxCas13d to generate higher performance.

| *Distance to Ideal Performance* | | |
|---|---|---|
| N-GID/GAI-C | s.s. | N-GAI/GID-C |
| 0.72 | **88** | 0.73 |
| 0.48 | **177** | 0.54 |
| 0.41 | **180** | 0.47 |
| 0.35 | **263** | 0.35 |
| 0.48 | **507** | 0.24 |
| 0.47 | **559** | 0.42 |
| 0.53 | **565** | 0.56 |
| 0.51 | **655** | 0.40 |
| 0.57 | **769** | 0.51 |
| 0.57 | **903** | 0.28 |
| ***Leakiness/Inducibility*** | | |
| 0.23/0.74 | | -0.16/0.82 |

**Supplementary Table 2. Characteristics of applied chemical inducible dimerization domains**

| Ligand | Origin | Toxicity in mammalian systems at working concentration | Potential endogenous target | Dimerization domain size (gene) | Dissociation constant (Kd) | Reversibility | Mechanism of action |
|---|---|---|---|---|---|---|---|
| Gibberellin | Plant hormone | no | no | 1.3kb | 1uM[3] | yes[3] | Dimerization |
| Abscisic acid | Plant hormone | no[4] | no | 1.6kb | 0.02uM-1.2uM[5] | yes[6] | Dimerization |
| Tamoxifen | Antiestrogen | no[7] | yes | 1kb | | hardly | Nuclear transportation |
| Danoprevir | Anti-viral drug | no[8] | no | 1.3kb | 0.02nM[9] | yes[9] | Dimerization |

**Supplementary Table 3. Split sites alignment among RfxCas13d, PspCas13b, and LwCas13a.**

| Cas13d s.s. | Cas13a s.s. | Cas13b s.s. |
|---|---|---|
| 149 | 397 | |
| 177 | 416 | 49 |
| 180 | 421 | |
| 263 | | 151 |
| 317 | 488 | |
| 456 | 605 | |
| 473 | 663 | |
| 507 | 786 | 250 |
| 559 | 826 | |
| 565 | 843 | 413 |
| 655 | 942 | 954 |
| 903 | | 1065 |

**Supplementary Table 4. Spacer sequences used in this study for mammalian RNA targeting.**

| Name | Sequence (5’ to 3’) |
|---|---|
| RfxCas13d B4GALNT1 pre-gRNA spacer[10] | CCTCCTGACCAGAAGCTGCCTGAAGGCTCA |
| RfxCas13d B4GALNT1 gRNA spacer[10] | CCTCCTGACCAGAAGCTGCCTG |
| RfxCas13d B2M pre-gRNA spacer[11] | ATATTAAAAAGCAAGCAAGCAGAATTTGGA |
| RfxCas13d B2M gRNA spacer[11] | ATATTAAAAAGCAAGCAAGCAGA |
| RfxCas13d CD46 pre-gRNA spacer[11] | AGACAATTGTGTCGCTGCCATCGAGGTAAA |
| RfxCas13d CD46 gRNA spacer[11] | AGACAATTGTGTCGCTGCCATCG |
| RfxCas13d mCh gRNA spacer[10] | ACCTTGAAGCGCATGAACTCCT |
| RfxCas13d non-targeting pre-gRNA 1[10] | TGCCACTACTGTTCATGATCAGGGCGATGG |
| PspCas13b mCh gRNA spacer[11] | ACCTTGAAGCGCATGAACTCCTTGATGATG |
| PspCas13b Cas13d gRNA spacer[11] | TTGTCATGTACACTTTGGAGCCGGACACGA |

**Supplementary Table 5. qPCR primer sequences used in this study for endogenous RNA targeting.**

| Transcript | Primer 1 (Forward, 5' to 3') | Primer 2 (Reverse, 5' to 3') | Amplicon size (bp) |
|---|---|---|---|
| GAPDH | TGACCTCAACTACATGGTTTACA | ATCGCCCCACTTGATTTTGG | 154 |
| B4GALNT | TGAGGCTGCTTTCACTATCCGC | GAGGAAGGTCTTGGTGGCAATC | 127 |
| B2M | CCACTGAAAAAGATGAGTATGCCT | CCAATCCAAATGCGGCATCTTCA | 126 |
| CD46 | TGGCTACCTGTCTCAGATGACG | GCATCTGATAACCAAACTCGTAAG | 115 |

**Supplementary Table 6. Timeline for the reversibility experiment.**

| **Day 0** | **Plate** | **Group a(+)** | **Group b** | **Group c** | **Group d** |
|---|---|---|---|---|---|
| Day1 | | 1 | + | | |
| Day2 | | 2 | 1 | + | |
| **Day3** | **sample and passage** | **3(-)** | **2** | **1** | **+** |
| Day4 | | -1 | 3(-) | 2 | 1 |
| Day5 | | -2 | -1 | 3(-) | 2 |
| **Day6** | **sample and passage** | **-3** | **-2** | **-1** | **3(-)** |
| Day7 | | -4 | -3 | -2 | -1 |
| Day8 | | -5 | -4 | -3 | -2 |
| **Day9** | **sample and passage** | **-6(+)** | **-5** | **-4** | **-3** |
| Day10 | | 1 | -6(+) | -5 | -4 |
| **Day11** | **Sample and passage** | **2** | **1** | **-6(+)** | **-5** |

*Numbers are the days of induction or recovery (-) a group of samples had experienced.*
*The symbol "(+)" represents addition of GA.*
*Blue colored cells represent samples during GA induction.*
*The symbol "(-)" represents removal of GA.*
*For detailed description, refer to the section: "Method Details- Stable Cell Line Generation and reversibility experiments"*
